# Platinum-Induced Ubiquitination of Phosphorylated H2AX by RING1A is Mediated by Replication Protein A in Ovarian Cancer

**DOI:** 10.1101/2020.04.30.070185

**Authors:** Shruthi Sriramkumar, Timothy D. Matthews, Ahmed H. Ghobashi, Samuel A. Miller, Pamela S. VanderVere-Carozza, Katherine S. Pawelczak, Kenneth P. Nephew, John J. Turchi, Heather M. O’Hagan

**Affiliations:** Cell, Molecular and Cancer Biology Graduate Program and Medical Sciences Program, Indiana University School of Medicine, Bloomington, IN, 47405, USA; Genome, Cell and Developmental Biology, Department of Biology, Indiana University Bloomington, Bloomington, IN, 47405, USA; Department of Medicine and Department of Biochemistry and Molecular Biology, Indiana University School of Medicine, Indianapolis, IN, 46202, USA; NERx Biosciences, Indianapolis, IN, 46201, USA; Indiana University Melvin and Bren Simon Cancer Center, Indianapolis, IN, 46202, USA; Department of Anatomy, Cell Biology and Physiology; Department of Obstetrics and Gynecology, Indiana University School of Medicine, Indianapolis, IN, USA; Department of Medical and Molecular Genetics, Indiana University School of Medicine, Indianapolis, IN, USA

**Author notes:** Corresponding author: Heather M. O’Hagan.

## Abstract

Platinum resistance is a common occurrence in high grade serous ovarian cancer (HGSOC) and a major cause of OC deaths. Platinum agents form DNA crosslinks, which activate nucleotide excision repair (NER), fanconi anemia (FA) and homologous recombination repair (HRR) pathways. Chromatin modifications occur in the vicinity of DNA damage and play an integral role in the DNA damage response (DDR). Chromatin modifiers, including polycomb repressive complex 1 (PRC1) members, and chromatin structure are frequently dysregulated in OC and can potentially contribute to platinum resistance. However, the role of chromatin modifiers in the repair of platinum DNA damage in OC is not well understood. We demonstrate that the PRC1 complex member RING1A mediates monoubiquitination of lysine 119 of phosphorylated H2AX (γH2AXub1) at sites of platinum DNA damage in OC cells. After platinum treatment, our results reveal that NER and HRR both contribute to RING1A localization and γH2AX monoubiquitination. Importantly, replication protein A (RPA), involved in both NER and HRR, mediates RING1A localization to sites of damage. Furthermore, RING1A deficiency impaired the activation of the G2/M DNA damage checkpoint and reduced the ability of OC cells to repair platinum DNA damage. Elucidating the role of RING1A in the DDR to platinum agents will allow for the identification of therapeutic targets to improve the response of OC to standard chemotherapy regimens.

## Introduction

Ovarian cancer (OC) is the most lethal gynecologic malignancy and fifth leading cause of death among women. Without the availability of adequate screening methods for early detection, the majority of patients are diagnosed with advanced stage disease (1). The standard of care for treatment of OC patients with advanced disease is surgical debulking followed by platinum-taxane based chemotherapy (2). High grade serous ovarian cancer (HGSOC), the most common OC histological type, initially responds to platinum-based therapy (3). However, up to 75% of responding patients relapse and eventually develop platinum-resistant disease. The survival rates of HGSOC have remained essentially unchanged for decades (3). Several mechanisms contribute to the development of platinum resistance, including increased drug efflux, decreased drug uptake, increased detoxification, increased DNA repair and reduced apoptotic response (4). Epigenetic mechanisms like histone modifications and promoter DNA methylation have also been associated with platinum resistance. Furthermore, dysregulation of chromatin modifiers in cancer leads to an altered DNA damage response (DDR) to chemotherapy agents through altered expression of genes involved in the DDR and altered repair of DNA lesions (5).

Platinum agents – cisplatin and carboplatin used for treatment of OC patients are DNA damaging agents. The cytotoxic activity of these agents is due to their ability to crosslink guanines. Cisplatin reacts with N7 positions of two guanines in the DNA forming intrastrand and interstrand crosslinks (ICLs) (6). Intrastrand adducts, the bulk of crosslinks/adducts formed by cisplatin, are repaired by the nucleotide excision repair (NER) pathway (7). Platinum ICLs are repaired by the Fanconi anemia (FA) and homologous recombination repair (HRR) pathways that result in activation of ataxia telangiectasia mutated (ATM) by auto-phosphorylation at S1981 (pATM) (8,9). Replication protein A (RPA)-coated single strand DNA (ssDNA) is a common structure formed during both NER and ICL repair that facilitates downstream DDR (10). ATM and ATR, which is activated by persistent single-stranded DNA, phosphorylate and activate downstream substrates including the histone variant H2AX.

Chromatin modifications occur in the vicinity of DNA damage, to promote signaling and repair of the damage by facilitating access to the DNA repair machinery (11). Crosstalk between DNA repair and chromatin has been explained by the “access, repair and restore” model in which the local chromatin around a site of DNA damage is modified to provide access to repair proteins, followed by repair of the damage and ultimately restoration of the chromatin to its original state (11). One such chromatin modification at sites of DNA damage is histone ubiquitination, including the monoubiquitination of lysine 119 of H2A/H2AX (H2A/H2AX K119ub1). RING domain-containing polycomb repressive complex 1 (PRC1) members RING1A/B in complex with BMI1 possess E3 ligase activity essential for adding the monoubiquitination mark on H2A/H2AX (12). BMI1 and RING1B localize to sites of IR and enzyme-induced double strand breaks (DSBs) and monoubiquitinate H2A/H2AX K119 (13–15). H2A/H2AX monoubiquitination also occurs in response to UV-induced DNA damage, facilitating recruitment of downstream repair proteins and repair activity (16,17). However, the role of chromatin modifiers and histone modifications in platinum DNA damage in OC remains poorly described. With increased repair of platinum adducts and altered DDR being common causes of platinum resistance, it is essential to understand histone modifications occurring at sites of platinum-induced DNA damage.

Here we demonstrate that RING1A, a member of the PRC1 complex, localizes to sites of platinum DNA damage and monoubiquitinates phosphorylated H2AX (γH2AX). We further show that both the global genome (GG)-NER and HRR pathways converge on γH2AXub1 and contribute to RING1A localization to the damage sites. Inhibition of the DNA binding of RPA decreases localization of RING1A to sites of cisplatin DNA damage. Furthermore, RING1A deficiency results in diminished activation of G2/M DNA damage checkpoint and reduced repair of cisplatin DNA damage. This is the first report of a role for RING1A in the platinum DNA damage response (DDR) in OC. Elucidating the role of RING1A in the DDR to platinum agents will allow for the identification of therapeutic targets to improve the response of OC to standard chemotherapy regimens.

## Materials and Methods

### Cell Culture

Cell lines used in the study were maintained at 37°C and 5% CO_2_. High grade serous ovarian cancer cells – OVCAR5 and Kuramochi were generously provided by Dr. Kenneth P. Nephew who had the lines authenticated by ATCC in 2018. OVCAR5 cells were cultured in DMEM 1X (Corning, PA # MT10013CV) with 10% FBS (Corning, PA #16000044) and Kuramochi cells were cultured in RPMI 1640 (Gibco, MA # MT10040CV) with 10% FBS without antibiotics as we have described previously (18). 293T cells obtained from ATCC were cultured in DMEM 1X with 10% FBS without antibiotics. All the cell lines used in the study were tested for mycoplasma using the Universal mycoplasma detection kit (ATCC, 30-1012K) on 10/10/2019. Cell lines used in all the experiments in the study were passaged for fewer than 15 passages. 154 mM NaCl (Macron Fine Chemicals #7581-12) solution in water was used to make the stock solution of cisplatin (Millipore Sigma, MA #232120). Stock solutions of carboplatin (Millipore Sigma, MA #216100) were made in water. Stock solutions of PRT4165 BMI1-RING1A E3 ligase inhibitor (Millipore Sigma, MA #203630) were made in DMSO. For experiments using this inhibitor, an equivalent amount of DMSO or inhibitor was added along with cisplatin and incubated for the 8 hours at 37°C in 5% CO_2_. Cells were pre-treated with DMSO or ATM inhibitor KU-55933 (Sigma Aldrich, MO #SML1109) for 1 hour prior to cisplatin or carboplatin treatment. Rad51 inhibitor B02 (Millipore Sigma, MA #553525) stock solutions were made in DMSO and pre-treated for 2 hours prior to cisplatin treatment. Stock solutions of RPA inhibitor NERx329 were made in DMSO. All the treatment doses and time points are specified in figure legends.

### Generation of stable knockdown lines using viral shRNAs

For knockdown of RING1A (Sigma, MO SHCLNG-NM_002931, TRCN0000021989, TRCN0000021990), BMI1 (Sigma, MO SHCLNG-NM_005180, TRCN0000020156, TRCN0000020157), XPC (Sigma, MO SHCLNG-NM_004628, TRCN0000083119), XPA (Sigma, MO SHCLNG-NM_000380, TRCN0000083196), CSB (Sigma, MO SHCLNG-NM_000124, TRCN0000436471) and empty vector TRC2 (Sigma, MO SHC201) lentiviral shRNA knockdown protocol from The RNAi Consortium Broad Institute was used. Briefly, 4 X 10^5^ 293T cells were plated on day 1 in DMEM 1X containing 10% FBS. On day 2, cells were transfected with shRNA of interest, EV control and packaging plasmids. Following transfection, 293T cells were incubated at 37°C and 5% CO_2_. On day 3, 16 -18 hours post transfection, media in the transfected flasks was replaced with fresh DMEM containing 10% FBS. Approximately 24 hours later, media was harvested and fresh DMEM + 10% FBS was added. The added media was harvested 24 hours later and pooled with media harvested on Day 4. This media harvested on day 4 and day 5 was filtered using 0.45 μm filter and concentrated using Spin-X concentrator (Corning, #431490).

### Antibodies

For western blot of endogenous proteins, anti-γH2AX (Cell Signaling Technology (CST), MA #9718, 1:1000), anti-Actin B (CST, MA #4970, 1:1000), anti-p-ATM S1981 (CST, MA #13050, 1:1000), anti-total ATM (CST, MA #2873, 1:1000), anti-total H2AX (CST, MA #2595, 1:1000), anti-RING1A (CST, MA #13069, 1:1000), anti-RING1B (Santa Cruz (SC), CA sc-101109, 1:1000), anti-H2AK119ub1 (CST, MA #8240,1:1000), anti-XPC (SC, CA sc-74410, 1:1000), anti-lamin B (SC, CA sc-6216, 1:1000; SC, CA sc-374015, 1:1000), anti-XPA (SC, CA sc-28353, 1:1000), anti-pRPA32 S33 (Bethyl Laboratories, TX A300-246A-M, 1:1000), anti-RPA32 (CST, MA #2208,1:1000), anti-phospho-Chk1 S345 (CST, MA #2348, 1:1000) and anti-total Chk1 (CST, MA #2360, 1:1000) antibodies were used. For immunofluorescence, anti-γH2AX (CST, MA #9718, 1:100), anti-RING1A (Abcam, CA ab175149, 1:100), anti-RPA32 (CST, MA #2208, 1:100), anti-Rad51 (Novus biological, CO NB100-148, 1:100), anti-H2AK119ub1 (Millipore Sigma, MA 05-678,1:100) and secondary Alexa Conjugate (CST, MA rat #4416, 1:1000, mouse #8890, 1:500, rabbit #4412, 1:1000 and, rabbit #8889,1:500) antibodies were used.

### Immunofluorescence with pre-extraction

OVCAR5 cells (2X10^5^) were cultured on coverslips in a 6-well plate and incubated at 37°C for 48 hours. After 48 hours, cells were either untreated or treated with cisplatin for 8 hours. For all ATM inhibitor experiments, OVCAR5 cells were not pretreated (Mock), pretreated with DMSO or 15 μM ATM inhibitor (ATMi) for 1 hour and then untreated (U) or treated with 12 μM cisplatin for 8 hours. For all Rad51 and RPA inhibitor experiments, cells were not pretreated (Mock) or pretreated with DMSO or 50 μM B02 (Rad51 inhibitor) or 8 μM NERx329 (RPAi) for 2 hours, respectively and then untreated (U) or treated with 12 μM cisplatin (T) for 8 hours. This was followed by pre-extraction using (0.5% Triton X-100 in 10 mM HEPES (pH7.4), 2 mM MgCl_2_, 100 mM KCl, 1 mM EDTA) and then fixed with 4% paraformaldehyde in PBS. Post fixation, cells were permeabilized using 0.5% Triton-X in PBS, blocked with 1% BSA in PBST (PBS + 0.1% Tween-20), incubated with appropriate primary antibodies as indicated and incubated with appropriate Alexa Fluor conjugated secondary antibodies. Coverslips were mounted using prolong gold antifade with DAPI (CST, MA #8961).

### Imaging and quantification

Images for all immunofluorescence experiments (except Supplementary Figure S1H) were acquired using the Leica SP8 scanning confocal system with the DMi8 inverted microscope. Leica LASX software (Leica Microsystems) was used for image acquisition. All the images were taken using 63X, 1.4NA oil immersion objective at RT. Images in Supplementary Figure S1H were obtained using Nikon NiE upright microscope with Hamamatsu Orca-Flash 2.8 sCMos high resolution camera. Following image acquisition, images were processed using Image J (National Institutes of Health, Bethesda, MD). For quantifying the percentage of cells with colocalization, at least 100 cells were scored. Each experiment was performed in 3 biological replicates. Co-localization was also confirmed using RGB profiler plugin on ImageJ as described previously (19).

### RNA isolation and quantitative reverse transcription PCR (RT-qPCR)

RNA extraction was performed using the RNAeasy mini kit (Qiagen, 74104). cDNA was synthesized using Maxima first strand cDNA synthesis kit for RT-qPCR (Thermo, MA K1642). FastStart Essential DNA green master (Roche, CA 06402712001) and *CSB* primers were used to amplify the cDNA. RT-qPCR primer sequences for CSB were *CSB*, forward, CTATGGTTGAGCTGAGGGCG and *CSB,* Reverse, GGGGATTCCCTCATTTGGCA

### Chromatin extraction

3 X 10^6^ cells were cultured in 150mm plates for approximately 48 hours. After UV treatment (see figure legend), cell pellets were used to perform nuclear extraction using CEBN (10 mM HEPES pH 7.8, 10 mM KCl, 1.5 mM MgCl_2_, 0.34 M sucrose, 10% glycerol, 0.2% NP-40, 1X protease inhibitor cocktail (Sigma, MO P5726), 1X phosphatase inhibitor (Thermo, MA 88266) and, N-ethylmaleimide (Acros organics, 128-53-0) and then washed with CEB buffer (CEBN buffer without NP-40) containing all the inhibitors. To extract the soluble nuclear fraction, after washing the cell pellets with CEBN buffer they were resuspended in soluble nuclear buffer (2 mM EDTA, 2 mM EGTA, all inhibitors) and rotated at 4°C for 30 min. The remaining cell pellet, i.e the total chromatin fraction, was lysed using 4 % SDS and analyzed by western blot.

### TCGA Analysis

Ovarian cancer patient datasets were compared to normal tissue using the TCGA TARGET GTEx dataset, accessed using Xenabrowser. Statistical significance was determined by pairwise comparisons using t-test with pooled standard deviations. P-values were adjusted for false discovery rates using Benjamini & Hochberg method.

### Statistical analysis

Percentage of cells with co-localization, relative densitometry and RT-qPCR data (presented as mean ± standard error (SEM)) were evaluated by using Student’s t-test in Graphpad prism and excel.

## Results

### RING1A contributes to platinum-induced monoubiquitination of γH2AX

As HGSOC is the most common OC histological type and most patients are treated with DNA damaging platinum agents (6), we utilized HGSOC cell lines as a model system to understand the role of chromatin modifiers in the DDR to platinum agents. We first determined the time point at which cisplatin induces H2AX phosphorylation in OC cells. γH2AX at S139 is a well-established marker of DNA breaks, including those that arise during the processing of platinum adducts by different repair pathways (20). Treatment of OVCAR5 cells with the IC50 dose of cisplatin caused a time-dependent increase in γH2AX (Figure 1A). Blotting for γH2AX with the same antibody previously used to also detect monoubiquitinated γH2AX (γH2AXub1) in response to DSB inducing agents (21) resulted in a band approximately 8 kD higher. γH2AX ub1 was first detected at the 8 hour time point and increased at the 16 hour time point when γH2AX levels were highest (Figure 1A). H_2_O_2_ induced comparable levels of γH2AX in OVCAR5 but monoubiquitination of γH2AX was not observed (Figure 1A), suggesting that γH2AXub1 was specifically induced by cisplatin and may play an important role in DDR to platinum agents. Cisplatin treatment also induced γH2AXub1 in HGSOC Kuramochi cells at 8 and 16 hour time points (Supplementary Figure S1A). In addition, treatment of OVCAR5 cells with the IC50 dose of carboplatin also induced γH2AXub1 after 16 and 24 hours (Supplementary Figure S1B). The increased time for detection of γH2AXub1 is in accordance with the finding that formation of DNA adducts by carboplatin were delayed compared to cisplatin due to differences in their aquation rates (22).

**Figure 1:**
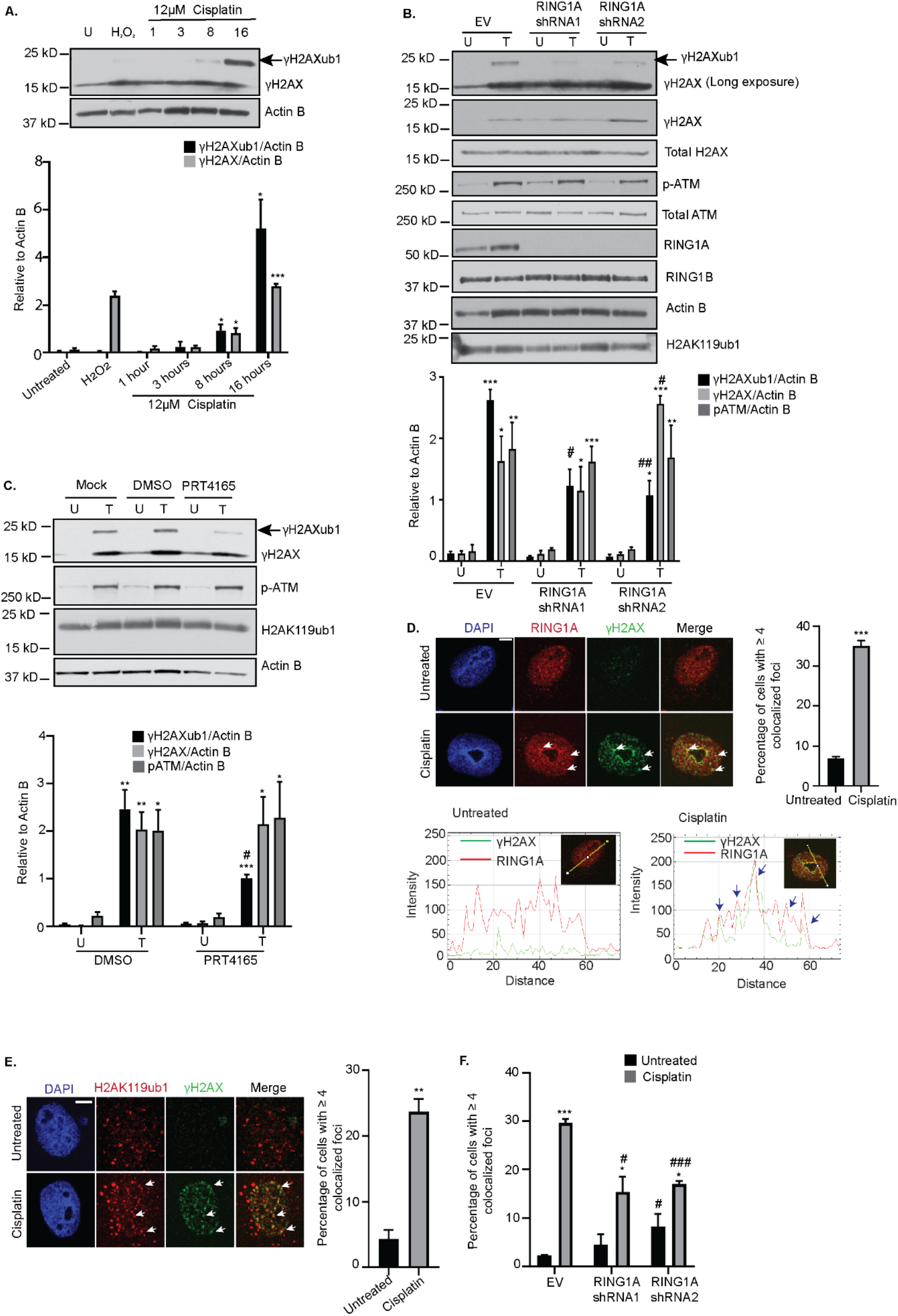
RING1A mediates platinum-induced monoubiquitination of γH2AX. **(A)** OVCAR5 cells were untreated (U) or treated with 12 μM cisplatin (IC50 dose) for 1, 3, 8 and 16 hours or with 2 mM H_2_O_2_ for 30 mins (used as a negative control). Cell lysates were analyzed by western blot. Graph depicts mean ± SEM of densitometric analysis of indicated proteins relative to Actin B at the indicated time points (N=3). **(B)** OVCAR5 cells infected with empty vector (EV) or 2 different RING1A shRNAs were untreated (U) or treated (T) with 12 μM cisplatin for 8 hours. Data is presented as in (A). OVCAR5 cells were either untreated (U) or treated with 12 μM cisplatin (T) alone or in combination with DMSO or 10 μM PRT4165 for 8 hours. Data is presented as in (A). OVCAR5 cells were treated 12 μM cisplatin for 8 hours. Immunofluorescence analysis was performed for RING1A (red) and the damage marker γH2AX (green). Merge image shows overlap of γH2AX and RING1A. White arrows indicate examples of RING1A foci that co-localize with γH2AX. Graph displays mean percentage of cells with ≥ 4 γH2AX and RING1A co-localized foci ± SEM (N=3). Scale bar = 5 μm. A representative RGB profile of untreated and cisplatin treated cell showing RING1A colocalization with γH2AX. Blue arrows point to foci which colocalize. **(E)** OVCAR5 cells were treated as in (D). Immunofluorescence analysis was performed for H2AK119ub1 (Red) and γH2AX (green). Merge image shows overlap of H2AK119ub1 and γH2AX. White arrows indicate examples of H2AK119 foci that co-localize with γH2AX. Graph displays mean percentage of cells with ≥ 4 γH2AX and H2AK119ub1 co-localized foci ± SEM (N=3). Scale bar = 5 μm. **(F)** OVCAR5 cells infected with empty vector (EV) or 2 different RING1A shRNAs were untreated or treated with 12 μM cisplatin for 8 hours and immunofluorescence was performed as in (E). Graph displays mean percentage of cells with ≥ 4 γH2AX and H2AK119ub1 co-localized foci ± SEM (N=3). Statistical significance was calculated using Student’s t-test. For all U versus T, P-values * < 0.05, ** <0.005, *** < 0.0005. For all EV versus RING1A KD or DMSO versus PRT4165, P-values, # < 0.05, ## < 0.005, ### < 0.005.

High expression of BMI1, RING1A and RING1B in recurrent ovarian tumors compared to primary tumors at presentation have been reported (23) and PRC1 members have been implicated in the repair of certain lesions. However, as the role of PRC1 complex members in the DDR to platinum agents in OC is not well understood, we first examined the expression of PRC1 members in TCGA OC patient data. Expression of BMI1 and RING1B was higher in in primary ovarian tumors compared to normal ovarian tissue (Supplementary Figure S1C). RING1A expression was higher than BMI1 or RING1B in normal ovarian tissue, but expression of RING1A in normal compared to primary tumor samples was not different (Supplementary Figure S1C). Based on these observations, we hypothesized that BMI1, RING1A/B contribute to platinum-induced γH2AXub1. To determine which of these PRC1 members contributed to platinum-induced ubiquitination, we independently knocked down BMI1, RING1A or RING1B. BMI1 or RING1B KD had no effect on cisplatin-induced γH2AXub1 (Supplementary Figure S1D and 1E). However, RING1A KD followed by cisplatin treatment reduced cisplatin-induced γH2AXub1 compared to an empty vector (EV) control (Figure 1B). The RING1A shRNAs had different effects on γH2AX levels. No change in γH2AX was observed after RING1A KD with shRNA1compared to a significant increase in γH2AX with shRNA2 (Figure 1B), suggesting that the decrease in cisplatin-induced γH2AXub1 observed with RING1A KD was not simply due to reduction in γH2AX levels. Double KD of RING1A and RING1B reduced cisplatin-induced γH2AXub1 to the same extent as RING1A KD alone, suggesting that RING1B does not contribute to the monoubiquitination of γH2AX (Supplementary Figure S1F). pATM and total ATM protein levels were not altered by RING1A KD plus cisplatin (Figure 1B), and basal H2AK119ub1 or RING1B levels were not significantly altered in RING1A KD cells (Figure 1B). RING1A KD also resulted in reduction in carboplatin-induced γH2AXub1 with no change in the levels of γH2AX (Supplementary Figure S1G).

The E3 ligase inhibitor PRT4165, inhibits BMI1-RING1A mediated ubiquitination of H2AK119 in a dose and time dependent manner and inhibits γH2AXub1 in response to IR-induced DSBs (24). Of the total H2A basally present in cells, 5 to 15% has been shown to be monoubiquitinated due to role of PRC1 in repression of homeobox (Hox) genes and X chromosome inactivation (25,26). Combined treatment with PRT4165 and cisplatin for 8 hours had no effect on basal H2AK119ub1 levels, though cisplatin-induced γH2AXub1 levels were reduced (Figure 1C). PRT4165 treatment had no effect on cisplatin-induced γH2AX and pATM levels (Figure 1C).

Proteins involved in DDR localize to and accumulate at DNA damage sites forming foci and BMI1 and RING1B form foci at sites of IR, enzyme-induced DSBs and sites of UV damage (13–15,27). Therefore, we were interested in examining if PRC1 members form foci at sites of cisplatin-induced DNA damage. Cisplatin treatment increased (P < 0.001) the number of cells with co-localization of RING1A and γH2AX foci (Figure 1D) demonstrating that RING1A is present at sites of DNA damage. RGB profiling of untreated and cisplatin treated cells demonstrated RING1A co-localization with γH2AX (Figure 1D), confirming the colocalization. Even though BMI1 KD had no effect on cisplatin-induced γH2AXub1 (Supplementary Figure S1D), BMI1 did form foci and localized to sites of platinum-induced DNA damage (Supplementary Figure S1H).

The role of RING1A in γH2AXub1 induction suggested lysine 119 as the site of monoubiquitination. However, an antibody against total H2AK119ub1, was unable to detect cisplatin-induced changes in H2Aub1 by western blot, likely because the basal PRC1-mediated H2AK119ub1 masks platinum-induced changes (in contrast, as shown in Figure 1C, the γH2AX antibody specifically detected cisplatin damage-induced γH2AX and γH2AXub1). Alternatively, using immunofluorescence, we observed that cisplatin treatment resulted in an increase in the percentage of cells with H2AK119ub1 at DNA damage foci (Figure 1E). A representative RGB profile of a cisplatin treated cell further confirmed co-localization (Supplementary Figure S2A). Treatment of RING1A KD cells with cisplatin reduced the percentage of cells with H2AK119ub1 at the damage foci compared to control, confirming that the platinum-induced monoubiquitination of γH2AX is mediated by RING1A and occurs on K119 (Figure 1F, Supplementary Figure S2B-D). Altogether, these results suggest that RING1A contributes to γH2AX monoubiquitination in response to platinum DNA damage.

### Knockdown of GG-NER proteins reduces cisplatin-induced γH2AXub1

Cisplatin primarily induces intrastrand adducts that can be repaired by both modes of the NER pathway (7,28), global genome NER (GG-NER repairs lesions throughout the genome) and transcription coupled NER (TC-NER repairs lesions recognized by stalling of RNA polymerase II (29)). Xeroderma pigmentosum complementation group C (XPC) is essential for GG-NER and the cockayne syndrome B (CSB) protein is involved in TC-NER (30,31). XPA has been implicated in early stages of both GG-NER and TC-NER (29,32). To determine which NER pathway plays a role in cisplatin-induced H2AX monoubiquitination, we knocked down NER pathway components. XPC KD decreased cisplatin-induced γH2AXub1, while CSB KD had no effect (Figure 2A, Supplementary Figure S3A-C). XPA KD also reduced cisplatin-induced γH2AXub1 (Figure 2B). Both XPC and XPA KD decreased γH2AX and pATM protein levels (Figure 2A, B), albeit these changes were not statistically significant.

**Figure 2:**
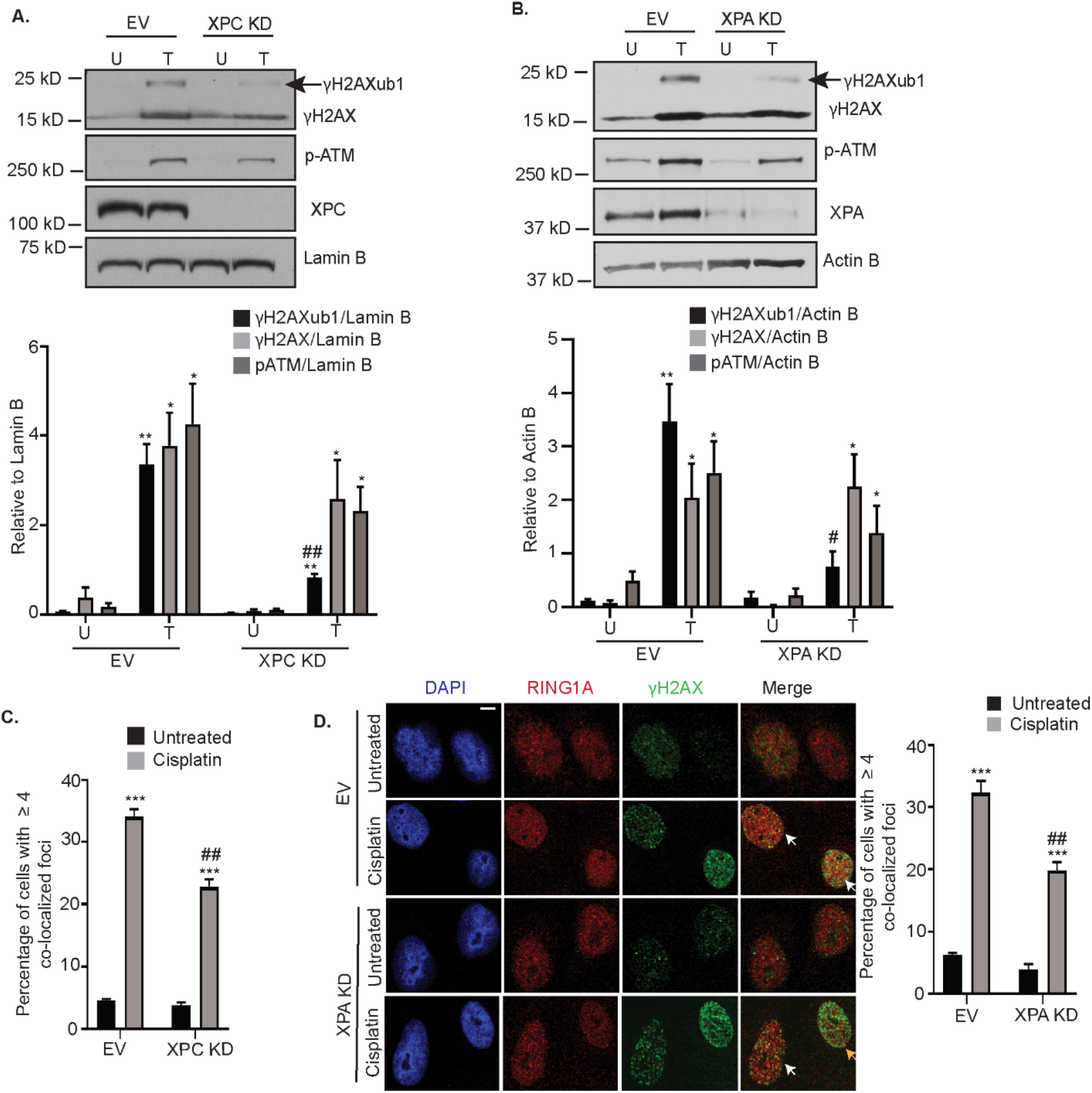
Knockdown of GG-NER proteins reduces cisplatin-induced γH2AXub1. Empty vector (EV) and XPC **(A)** or XPA **(B)** KD OVCAR5 cells were untreated (U) or treated with 12 μM cisplatin for 8 hours (T). Cell lysates were analyzed by western blot. Graphs depict mean ± SEM of densitometric analysis of indicated proteins relative to indicated housekeeping genes (N=3). **(C)** OVCAR5 EV and XPC KD cells were treated with 12 μM cisplatin for 8 hours. Immunofluorescence analysis was performed for RING1A (red) and the damage marker γH2AX (green). Graph depicts mean percentage of cells having ≥ 4 γH2AX and RING1A co-localized foci ± SEM (N=3). **(D)** OVCAR5 EV and XPA KD cells were treated and analyzed as in (C). White arrows indicate examples of cells showing γH2AX and RING1A colocalization while yellow arrows indicate examples of cells which do not have γH2AX and RING1A co-localization (yellow arrow indicates reduction in yellow foci). Scale bar = 5 μm. Statistical significance was calculated using Student’s t-test. For all U versus T, P-values * < 0.05, ** <0.005, *** < 0.0005. For all EV versus XPC or XPA KD, P-values # < 0.05, ## < 0.005, ### < 0.005.

We next examined the effect of XPC and XPA KD on RING1A localization to sites of cisplatin DNA damage. Both XPC and XPA KD decreased the percentage of cells showing RING1A localization to DNA damage foci (Figure 2C, D, Supplementary Figure S3D), suggesting that GG-NER contributes to RING1A localization to sites of platinum DNA damage. However, RING1A localization was not completely abrogated by XPC or XPA KD suggesting that other repair pathways may also contribute to RING1A localization to DNA damage sites.

### Inhibition of HRR reduces platinum-induced γH2AXub1

ICLs formed by cisplatin block DNA replication and transcription and are processed by the FA and HRR pathways (7). As expected, treatment of OVCAR5 or Kuramochi cells with the IC50 doses of cisplatin resulted in ATM phosphorylation at S1981 (Figure 3A, Supplementary Figure S1A), and carboplatin also activated pATM in OVCAR5 cells (Supplementary Figure S1B). Accumulation of H2Aub1 at sites of enzyme-induced DSBs was shown to be dependent on ATM (33), and we hypothesized that ATM inhibition would alter platinum-induced γH2AXub1. Treatment of OVCAR5 cells with ATMi (15 μM KU-55933) followed by 8 hours cisplatin treatment reduced cisplatin-induced γH2AXub1 (Figure 3B). ATMi also reduced cisplatin-induced γH2AXub1 in Kuramochi cells and carboplatin-induced γH2AXub1 in OVCAR5 cells (Supplementary Figure S4A, B). Furthermore, treatment with KU-55933 followed by cisplatin reduced RING1A localization to DNA damage foci in comparison to the vehicle control (Figure 3C, Supplementary Figure S4C). As KD or inhibition of RING1A had no effect on platinum-induced ATM phosphorylation at S1981 or its activation (Figure 1B and C), we suggest that RING1A functions downstream of pATM during repair of platinum DNA damage.

**Figure 3:**
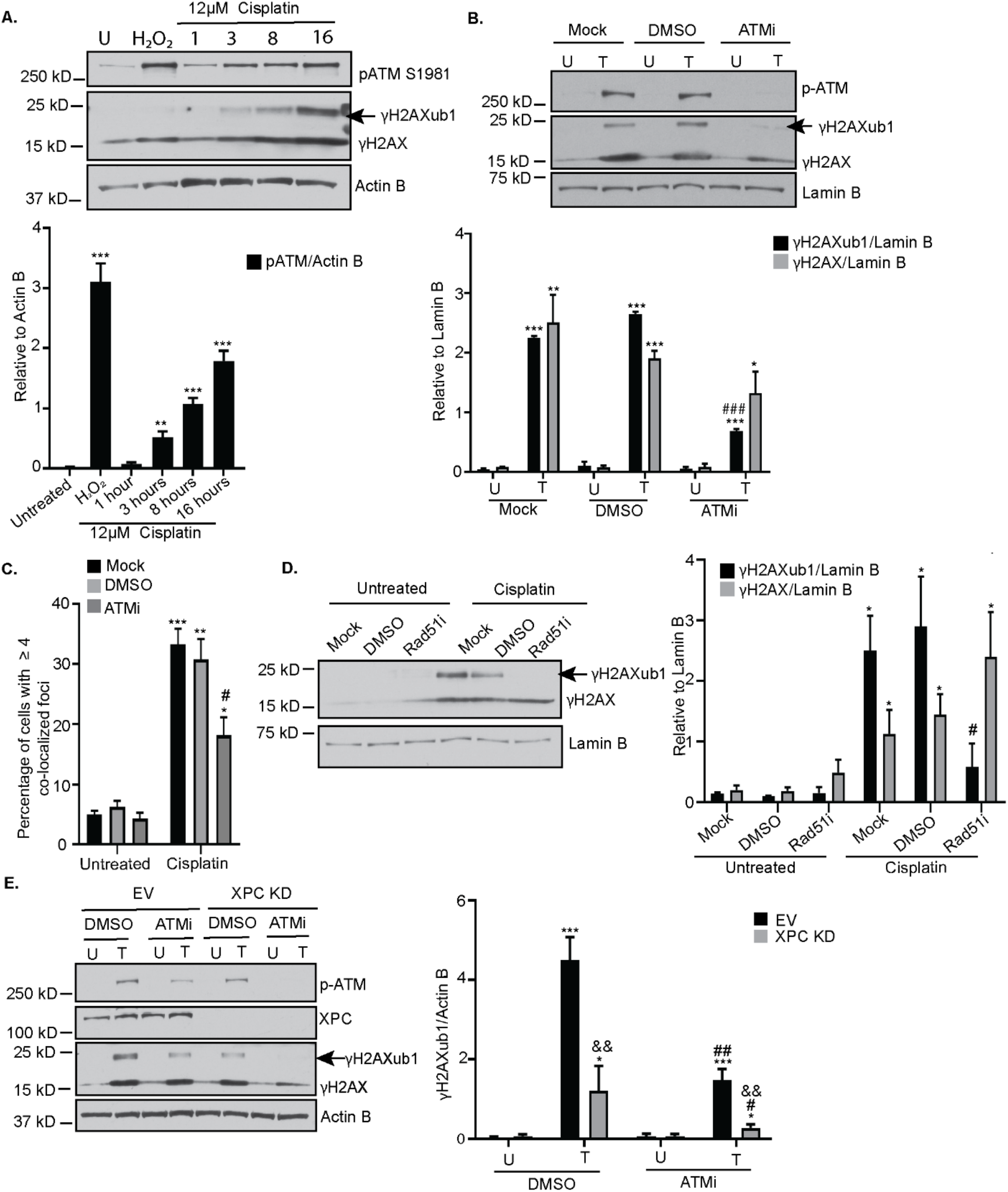
Inhibition of HRR reduces platinum-induced γH2AXub1. **(A)** OVCAR5 cells lysates used in Figure 1A were blotted for pATM. Graphs depict mean ± SEM of densitometric analysis of indicated proteins relative to the indicated house-keeping gene (N=3). **(B)** OVCAR5 cells were not pretreated (mock) or pretreated with either DMSO or 15 μM ATM inhibitor (ATMi) Ku-55933 for 1 hour and then untreated (U) or treated (T) with 12 μM cisplatin for 8 hours. Data is presented as in (A). **(C)** OVCAR5 cells were treated as in (B). Immunofluorescence analysis was performed. Data represents percentage of cells having ≥ 4 co-localized γH2AX and RING1A foci ± SEM (N=3). **(D)** OVCAR5 cells were not pretreated (mock) or pretreated with DMSO or 50 μM Rad51 inhibitor (B02) for 2 hours and then untreated (U) or treated (T) with 12 μM cisplatin for 8 hours. Data is presented as in (A). **(E)** EV or XPC KD OVCAR5 cells were either treated with DMSO or ATM inhibitor (KU-55933) alone (U) or treated (T) with cisplatin for 8 hours. Data is presented as in (A). Statistical significance was calculated using Student’s t-test. For all U versus T, P-values * < 0.05, ** <0.005, *** < 0.0005. For all mock or DMSO versus ATMi or Rad51i, P - values # < 0.05, ## < 0.005, ### < 0.005. For EV versus XPC KD in figure 3E, P-values & < 0.05, && < 0.005, &&& < 0.0005.

An important step in HRR is homology search and strand exchange, catalyzed by Rad51 (34). In response to cisplatin treatment, an expected increase in the percentage of cells having co-localization of Rad51 with γH2AX foci was observed (Supplementary Figure S4D). Inhibition of Rad51 using B02 followed by cisplatin treatment also reduced γH2AXub1 without altering γH2AX levels (Figure 3D). As a positive control for efficacy of B02 treatment, OVCAR5 cells treated with B02 had a significant reduction in Rad51 foci formation in response to IR (Supplementary Figure S4E) as previously shown by Huang F et al (35). Overall, these results demonstrate that HRR contributes to RING1A localization to sites of DNA damage and platinum-induced γH2AXub1.

Having demonstrated that both GG-NER and HRR contribute to platinum-induced γH2AXub1, we next sought to investigate the combined effect of ATMi and XPC KD on cisplatin-induced γH2AXub1. Cisplatin treatment of EV OVCAR5 cells treated with ATM inhibitor or KD of XPC alone resulted in a reduction in cisplatin-induced γH2AXub1 (Figure 3E), consistent with our previous results (Figure 3B and 2A). ATMi plus cisplatin treatment further reduced the level of γH2AXub1 in XPC KD cells (Figure 3E), indicating that the two pathways function in parallel in response to cisplatin DNA damage and both result in γH2AXub1.

### RPA facilitates RING1A localization to sites of cisplatin-induced DNA damage

Based on our data that both GG-NER and HRR pathways result in RING1A-mediated γH2AXub1, we hypothesized that a protein involved in both pathways facilitates RING1A localization to sites of platinum DNA damage. RPA has been implicated as a key player in both GG-NER and DSB repair pathways HRR and non-homologous end joining (NHEJ) (36,37). The RPA32 subunit of RPA is known to be phosphorylated by ATR and ATM in response to replication stress and IR-induced DSBs regulating downstream protein-protein and protein-DNA interactions (38,39). RPA32 Serine 33 (S33) is phosphorylated in response to crosslinking agents like UV (37). Cisplatin but not H_2_O_2_ induced RPA32 phosphorylation at S33 (Figure 4A). Cisplatin treatment increased RPA32 punctate foci which co-localized with the damage marker γH2AX compared to untreated cells (Figure 4B, Supplementary Figure S5A) as has been demonstrated by others (40). Furthermore, cisplatin treatment increased the percentage of cells with RPA32 foci that co-localized with RING1A in comparison to untreated cells (Figure 4C, Supplementary Figure S5B).

**Figure 4:**
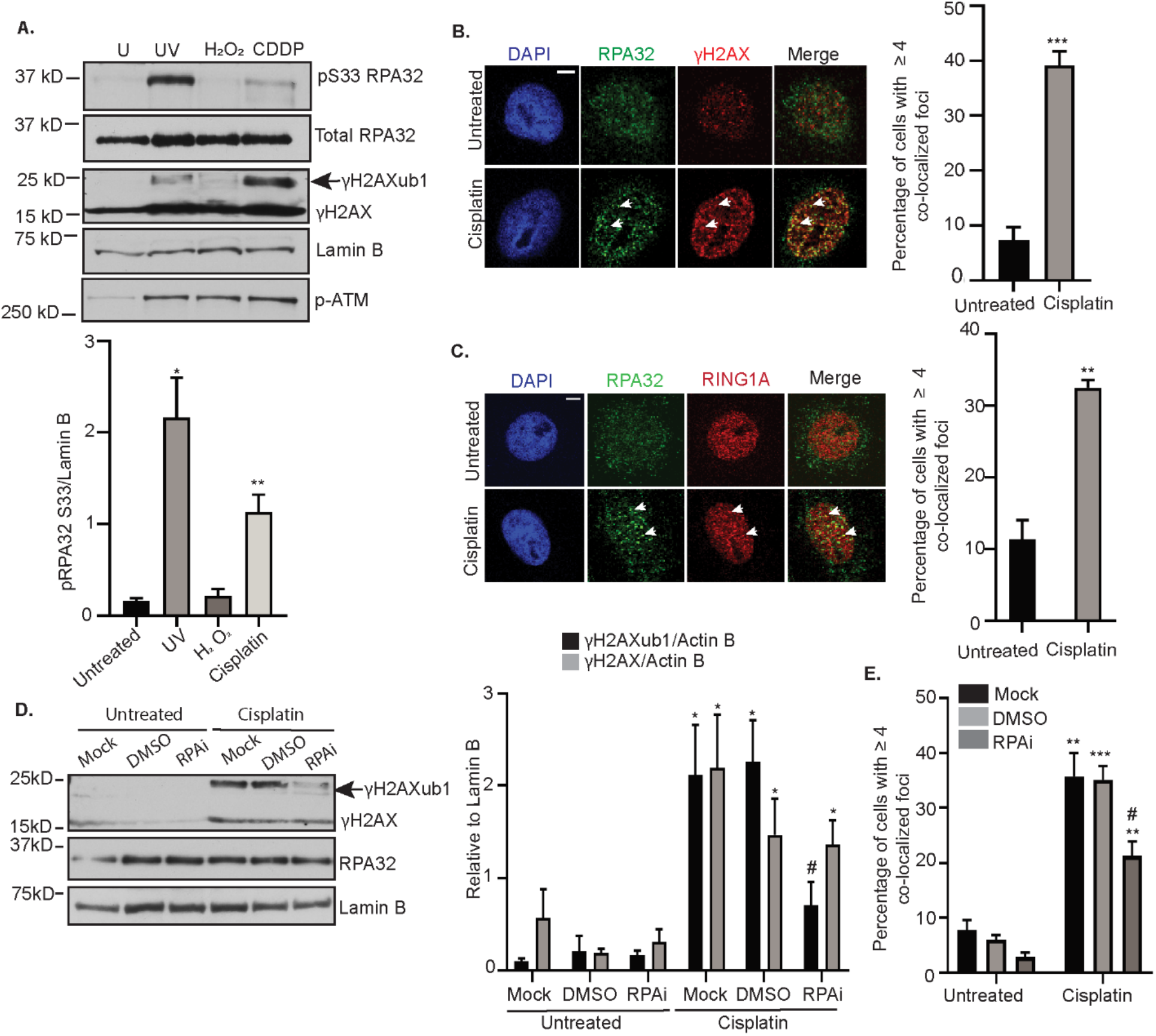
RPA facilitates RING1A localization to sites of cisplatin-induced DNA damage. **(A)** OVCAR5 cells were untreated (U) or treated with 200 mJ/cm^2^ UV (positive control) followed by 15 mins recovery, 2 mM H_2_O_2_ for 30 minutes, or 12 μM cisplatin (CDDP) for 8 hours. H_2_O_2_ and UV were negative and positive controls, respectively for pS33 RPA32 phosphorylation. Graphs depict mean ± SEM densitometric analysis of indicated proteins relative to the indicated house-keeping gene (N=3). **(B&C)** OVCAR5 cells were treated with 12 μM cisplatin for 8 hours, followed by immunofluorescence analysis for indicated antibodies. Graph depicts mean ± SEM percentage of cells with ≥4 RPA32 and γH2AX **(B)**, and RPA32 and RING1A **(C)** co-localized foci of N=3 biological replicates. White arrows indicated examples of γH2AX and RING1A foci which co-localize. Scale bar = 5 μm. **(D)** OVCAR5 cells were not pre-treated (Mock) or pre-treated with DMSO or 8 μM NERx329 (RPAi) for 2 hours followed by 8 hours of cisplatin treatment. Data is presented as in (A). **(E)** OVCAR5 cells were not pre-treated (Mock) or pre-treated with DMSO or 8 μM RPAi for 2 hours and then untreated or treated with cisplatin for 8 hours. Immunofluorescence was performed and analyzed as in (B). Statistical significance was calculated using Student’s t-test. For all U versus T, P-values * < 0.05, ** <0.005, *** < 0.0005, ns – not significant. For all mock or DMSO versus RPAi, P - values # < 0.05, ## < 0.005, ### < 0.005.

To test the hypothesis that RPA mediates the localization of RING1A to sites of platinum DNA damage, we pre-treated OVCAR5 cells with a reversible RPA inhibitor (RPAi), NERx329, which inhibits RPA binding to single stranded DNA (41,42). RPAi reduced cisplatin-induced γH2AXub1 without altering γH2AX levels (Figure 4D). Demonstrating the efficacy of RPAi, OVCAR5 cells pretreated with RPAi prior to UV treatment had a significant reduction in the UV-induced binding of RPA32 to chromatin in comparison to the control (Supplementary Figure S5C). RPAi also reduced the percentage of cells with co-localization of RING1A and γH2AX foci (Figure 4E, Supplementary Figure S5D). Altogether, this data suggests that RPA facilitates RING1A localization to sites of cisplatin DNA damage, resulting in γH2AXub1.

### RING1A contributes to the repair of cisplatin-induced DNA damage

It was next of interest to investigate the effect of RING1A KD on events downstream of RPA binding to ssDNA, a step which occurs in both GG-NER and HRR pathways (10,43). RPA binding to ssDNA recruits the ATR-ATRIP complex to damage sites which further results in phosphorylation of RPA32 at S33 and checkpoint kinase 1 (Chk1) at S345, which is required for the activation of G2/M DNA damage checkpoint (43–46). RING1A KD resulted in reduced platinum-induced phosphorylation of RPA32 at S33 and Chk1 at S345. (Figure 5A-C; validated using a second RING1A shRNA, RING1A shRNA2; Supplementary Figure S6A). To examine the effect of RING1A KD on the ability of OC cells to repair cisplatin DNA damage, we treated OVCAR5 cells with 6 μM cisplatin followed by recovery in platinum-free media. The percentage of cells with γH2AX foci, a measure of the ability of cells to repair DNA damage (20,47), was maximal at 24 hours of recovery in both EV and RING1A KD cells (Figure 5D, E). At 72 hours of recovery, the percentage of γH2AX positive cells observed in EV cells decreased to approximately 38% relative to the 48 hour time point, suggesting repair of the DNA damage. In contrast, the percentage of RING1A KD cells with γH2AX foci was unchanged at 72 and 96 hours post recovery, with approximately 65% and 62% RING1A KD cells having persistent γH2AX foci, respectively (Figure 5D, 5E; validated by using RING1A shRNA2; Supplementary Figure S6B).

**Figure 5:**
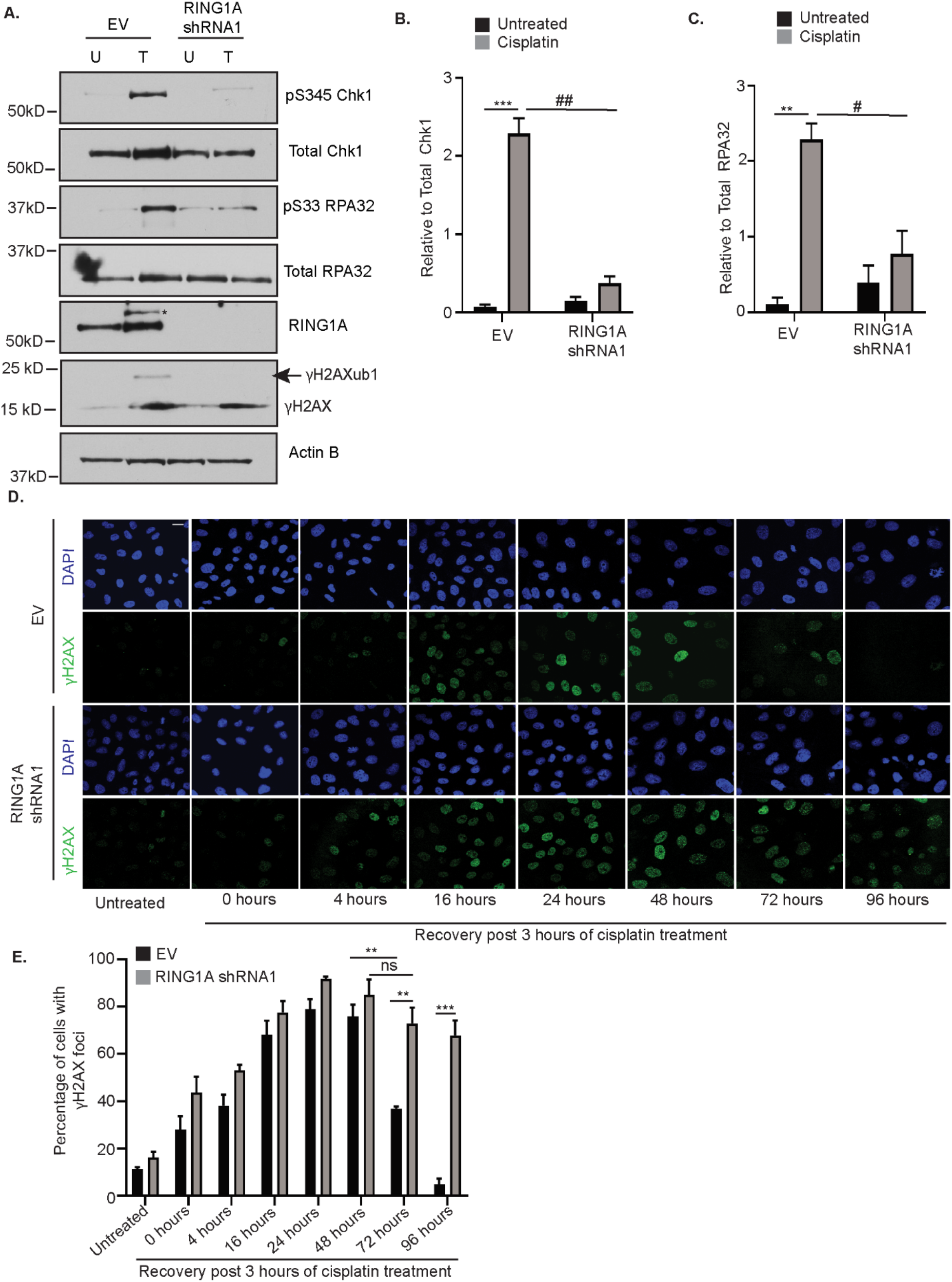
RING1A contributes to the repair of cisplatin-induced DNA damage. **(A)** OVCAR5 cells infected with EV or RING1A shRNA1 were untreated (U) or treated with 12μM cisplatin for 8 hours. Cell lysates were analyzed by western blot.* denotes non-specific band. Graph depicts mean ± SEM of densitometric analysis of N=3 biological replicates of pS345 Chk1 relative to total Chk1 **(B)** and pS33 RPA32 relative to total RPA32 **(C)**. For all U versus T, P-values * < 0.05, ** <0.005, *** < 0.0005, ns - not significant. For all EV versus RING1A KD, P - values # < 0.05, ## < 0.005, ### < 0.005. **(D)** EV or RING1A KD OVCAR5 cells were treated with 6 μM cisplatin for 3 hours. Cells were then allowed to recover in platinum-free media for the indicated time points followed by immunofluorescence for γH2AX (green) as a surrogate for DNA damage. Representative images of OVCAR5 EV and RING1A KD cells at the indicated time points are displayed. Scale bar = 20 μm. **(E)** Graph depicts mean percentage of cells with γH2AX foci ± SEM in N=3 biological replicates. For all EV versus RING1A KD, P-values * < 0.05, ** <0.005, *** < 0.0005, ns – not significant.

## Discussion

Aberrant DNA damage signaling and repair are prominent features of ovarian tumorigenesis and platinum resistance (48). Furthermore, chromatin modifiers are also dysregulated in OC and have been targeted to improve response to chemotherapy agents. While chromatin modifications have been implicated in the DDR to several types of DNA damaging agents, their role in the repair of DNA damage caused by platinum-based chemotherapy agents has not been well-studied. Here we demonstrate for the first time a role for the PRC1 member RING1A and monoubiquitination of γH2AX in the DDR to platinum-induced DNA damage in HGSOC.

Platinum agents are a treatment mainstay for OC. DNA damage induced by platinum agents are repaired by NER, FA and HRR pathways (7,49,50). Here, we have implicated both GG-NER and HRR pathways in localization of RING1A to sites of platinum DNA damage and the subsequent increase in γH2AXub1 (Supplementary Figure S6C). We speculate that platinum-induced γH2AXub1 occurs due to double and single stranded DNA break intermediates which are generated during the processing of cisplatin intra- and interstrand crosslinks (Supplementary Figure S6C). The involvement of multiple repair pathways in facilitating RING1A mediated repair of platinum adducts supports the idea that anticancer therapies targeting multiple DNA repair hubs should be exploited in OC and other cancer types as suggested by Deitlein et al.(51).

Exhaustion of RPA (a common player in both GG-NER and HRR) during replication stress is demonstrated to be a vital determinant of cisplatin resistance in HGSOC cells (52). We demonstrate that RPA mediates RING1A localization to damage sites connecting NER and HRR pathways to RING1A mediated γH2AXub1 (Supplementary Figure S6C). Our findings are consistent with the idea that RPA can orchestrate the localization of many proteins to sites of DNA damage to promote repair (53,54). Future work is needed to further understand the mechanism that connects RING1A to RPA. The RPA inhibitor used in this study blocks the binding of RPA’s OB folds to DNA (42). An inhibitor targeting RPA-DNA interaction such as NERx329 has advantages over other types of RPA inhibitors because it inhibits RPA independent from its phosphorylation status. RPA plays a vital role in DNA replication and other DNA repair pathways in addition to NER and HRR. Developing RPA inhibitors like NERx329 to be used in combination with platinum could be beneficial in killing rapidly proliferating cancer cells and repair proficient chemoresistant OC cells.

Recurrent ovarian tumors have high expression of PRC1 members – BMI1, RING1A and RING1B (23). We demonstrate that RING1A contributes to platinum-induced monoubiuquitination of γH2AX. Because of limitations of commercially available H2AK119ub antibodies we detected ubiquitination using an antibody against γH2AX. Our data suggests that cisplatin-induced monoubiquitination of H2AX occurs after its phosphorylation. Although we focused on RING1A-mediated γH2AXub1, monoubiquitination is also likely occurring on H2A in response to platinum. It is possible that sites where damage occurred already contained monoubiquinated H2AX. However, we do not believe this is the case as phosphorylation of H2AX is also induced by H_2_O_2_ without detection of monoubiquitinated γH2AX (see Figure 1A and 3A). The persistence of γH2AX foci in RING1A KD cells indicates that RING1A promotes the repair of platinum DNA damage. While RING1A KD may alter chromatin dynamics and hence indirectly result in persistent γH2AX foci, the alteration of the G2/M checkpoint suggests that RING1A KD alters the DDR. Together, our data demonstrates that RING1A plays an important role in repair of cisplatin-induced DNA damage in OC cells.

PRC1 complexes are heterogenous in nature and can contain several different E3 ligases (55). In our study, BMI1 localized to sites of platinum DNA damage (Supplementary Figure S1H), however, KD of BMI1 or RING1B did not affect platinum-induced γH2AXub1 and the presence of RING1B did not compensate for the depletion of RING1A in regard to platinum-induced γH2AXub1 (Supplementary Figure S1D, S1E). We reason that as there are multiple PRC1 and PRC1-like complexes, and RING1A and RING1B can replace each other, it is likely that a PRC1 or PRC1-like complex having predominantly RING1A as the E3 ligase is involved in the DDR to platinum agents in OC cells. Further studies are essential to determine which PRC1 or PRC1-like complex members in addition to RING1A are involved in DDR to platinum agents in other cancer types and why specifically RING1A but not RING1B is involved in DDR of platinum lesions in OC cells.

The main obstacle in the use of platinum agents for treatment of OC and other cancers like colon and lung is the development of chemotherapy resistance. Our study has furthered our understanding of the role of RING1A in the DDR to platinum agents in OC and has established a link between repair pathways and localization of RING1A to damage sites. RPA inhibitors like the one used in our study have shown efficacy in regressing the growth of tumors in xenograft model of non-small cell lung carcinoma (41). Evaluating the effect of such inhibitors in OC mouse models will aid in improving response of recurrent OC patients to standard chemotherapy regimens. Furthermore, the activities of many chromatin modifiers are druggable and so understanding their role in DDR in OC cells can lead to potential therapeutic targets. As platinum-based agents are used in chemotherapy regimens in several other cancers our findings warrant further investigation of the role of PRC1 in the repair of platinum lesions in other cancer types as well.

## Acknowledgements

SS dedicates this work in the loving memory of her mother Brinda Sriramkumar. This research was funded by Ovarian Cancer Research Alliance grant number 458788 to HMOH and KPN. SS was supported by the Doane and Eunice Dahl Wright Fellowship generously provided by Ms. Imogen Dahl. We thank the Indiana University Light Microscopy Imaging Center for their assistance.

## Author Contributions

HMOH and SS conceived and designed all the experiments and wrote the manuscript. TDM contributed Fig.5D and Supp. Fig.6B. AHG contributed Supp. Fig.1B, 1G and Supp. Fig.4B. SAM contributed Supp. Fig. 1C. SS performed and analyzed all the other experiments. PSVC, KSP and JJT provided RPA inhibitor NERx329. HMOH, SS, KPN and JJT edited and proofread the manuscript.

