## Supplemental Figures for "Platinum-Induced Ubiquitination of Phosphorylated H2AX by RING1A is Mediated by Replication Protein A in Ovarian Cancer"

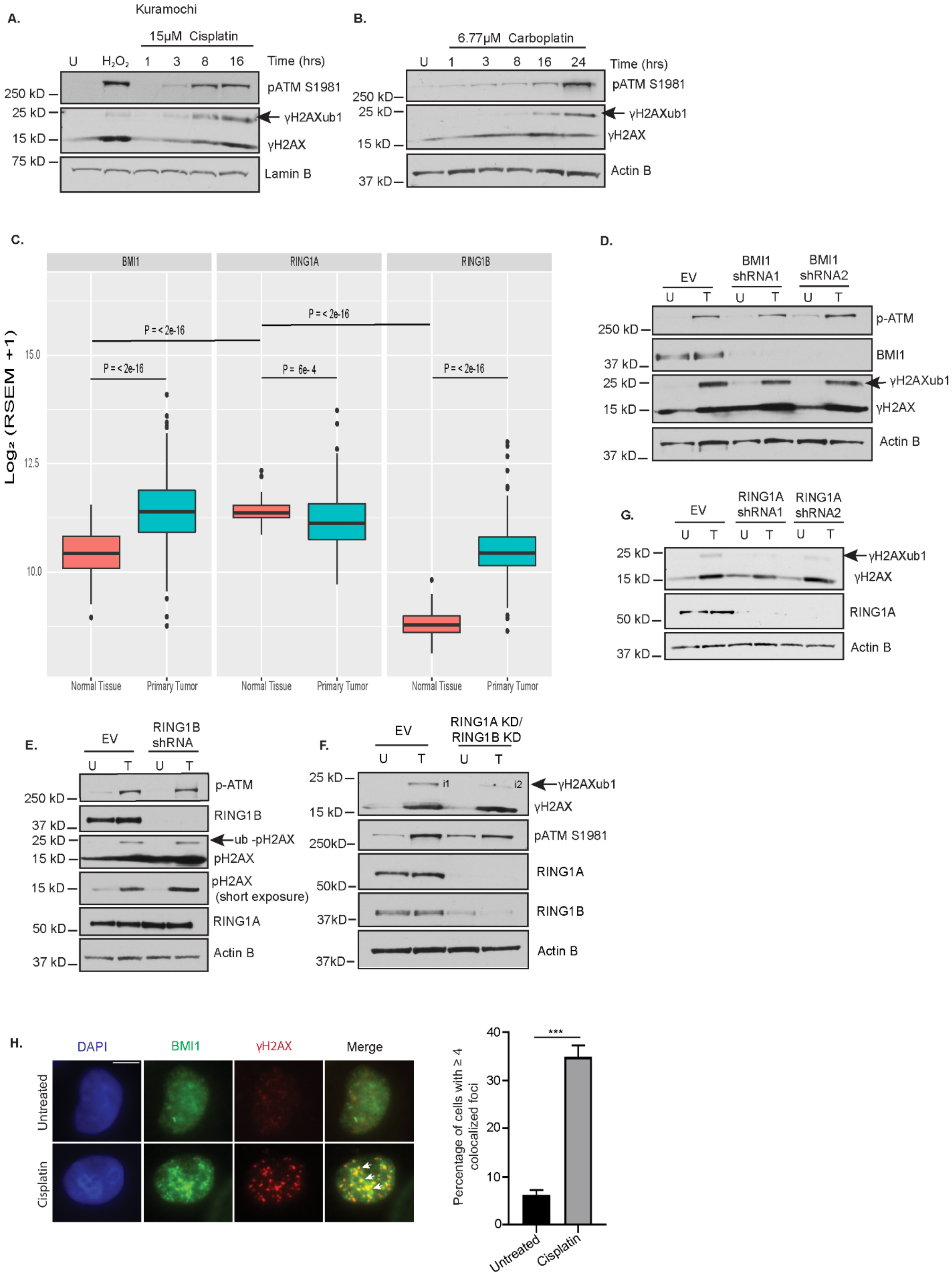
A.
Kuramochi
15μM Cisplatin
U H2O2 1 3 8 16 Time (hrs)
pATM S1981
γH2AXub1
γH2AX
Lamin B
250 kD
25 kD
15 kD
75 kD
B.
6.77μM Carboplatin
U 1 3 8 16 24 Time (hrs)
pATM S1981
γH2AXub1
γH2AX
Actin B
250 kD
25 kD
15 kD
37 kD
C.
BMI1
RING1A
RING1B
Log2 (RSEM +1)
P = < 2e-16
P = 6e-4
Normal Tissue
Primary Tumor
D.
EV
BMI1 shRNA1
BMI1 shRNA2
U T U T U T
p-ATM
BMI1
γH2AXub1
γH2AX
Actin B
G.
EV
RING1A shRNA1
RING1A shRNA2
RING1A
50 kD
E.
RING1B shRNA
p-ATM
RING1B
ub -pH2AX
pH2AX
pH2AX (short exposure)
F.
RING1A KD/ RING1B KD
H.
DAPI
BMI1
γH2AX
Merge
Untreated
Cisplatin
Percentage of cells with ≥ 4 colocalized foci
***

**Supplementary Figure S1: RING1A mediates monoubiquitination of γH2AX in response to carboplatin**

**(A)** Kuramochi cells were untreated (U), treated with 2 mM $H_2O_2$ for 30 minutes or 15 μM cisplatin (IC50 dose) for the indicated time points. Lysates were analyzed by WB (N=3). **(B)** OVCAR5 cells were untreated (U) or treated with 6.77 μM carboplatin (IC50 dose) for the indicated time points. Lysates were analyzed by WB (N=3). **(C)** Box plots of BMI1, RING1A and RING1B expression in normal ovary from GTEX dataset and primary tumor samples from TCGA. OVCAR5 cells were infected with empty vector (EV) or 2 independent BMI1 viral shRNAs **(D)** or RING1B viral shRNA **(E)** or RING1A and RING1B viral shRNA **(F)** and then untreated (U) or treated with 12 μM cisplatin (T) for 8 hours. Lysates were analyzed by WB (N=3). Intensity 1 (i1) is 2.40 and intensity 2 (i2) is 1.49 relative to respective loading control. **(G)** OVCAR5 cells were infected with EV or 2 independent RING1A viral shRNAs and then untreated (U) or treated with 6.77 μM carboplatin (IC50 dose) for 24 hours. Cell lysates were analyzed by WB. **(H)** Kuramochi cells were treated with 15 μM cisplatin for 8 hours and then immunofluorescence analysis was performed for BMI1 (green) and the damage marker γH2AX (red). Merge image shows overlap of γH2AX and BMI1. White arrows indicate examples of BMI1 foci that co-localize with γH2AX. Graph displays mean percentage of cells with ≥ 4 γH2AX and BMI1 co-localized foci ± SEM (N=3). Scale bar = 5 μm.

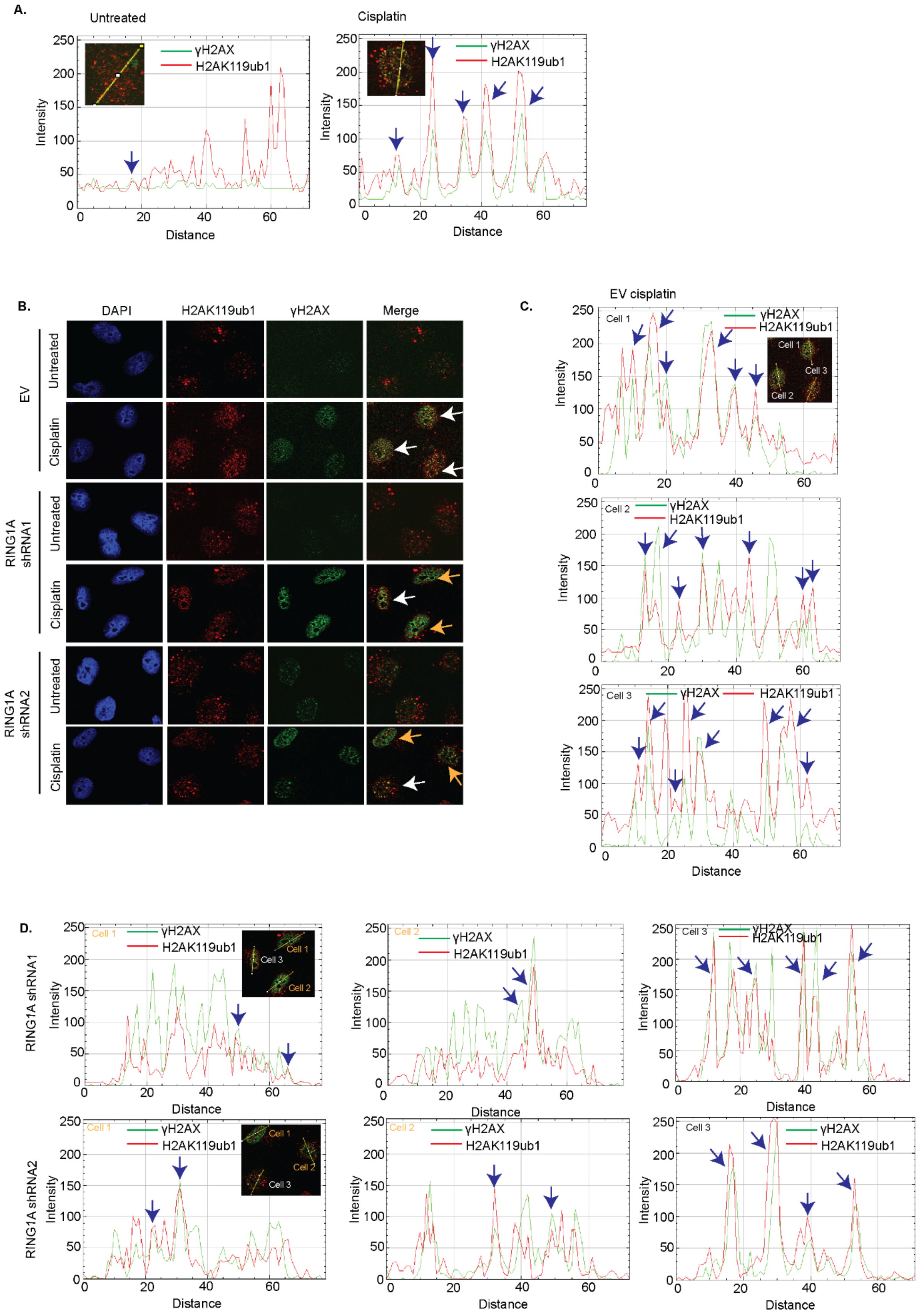
A.
Untreated
Cisplatin
γH2AX
H2AK119ub1
Intensity
Distance
B.
DAPI
H2AK119ub1
γH2AX
Merge
EV
Untreated
Cisplatin
RING1A shRNA1
RING1A shRNA2
C.
EV cisplatin
Cell 1
Cell 2
Cell 3
D.
RING1A shRNA1
RING1A shRNA2

**Supplementary Figure S2: RING1A monoubiquitinates γH2AX at lysine 119 in response to cisplatin treatment**

**(A)** A representative RGB profile of an untreated and cisplatin treated cell from Figure 1E showing H2AK119ub1 colocalization with γH2AX. Blue arrows point to foci which colocalize. **(B)** Representative images of untreated and cisplatin treated OVCAR5 EV and RING1A KD cells showing H2AK119ub1 colocalization with γH2AX. White arrows point to cells showing colocalization (Note the yellow colored foci in EV cisplatin treated cells) and yellow arrows point to cells with reduced colocalization (Note the decrease of yellow foci in RING1A KD cisplatin treated cells). Representative RGB profiles of EV cisplatin treated cells **(C)** and RING1A KD cisplatin treated cells **(D)** showing colocalization of γH2AX and H2AK119ub1. Blue arrows point to colocalizing foci.

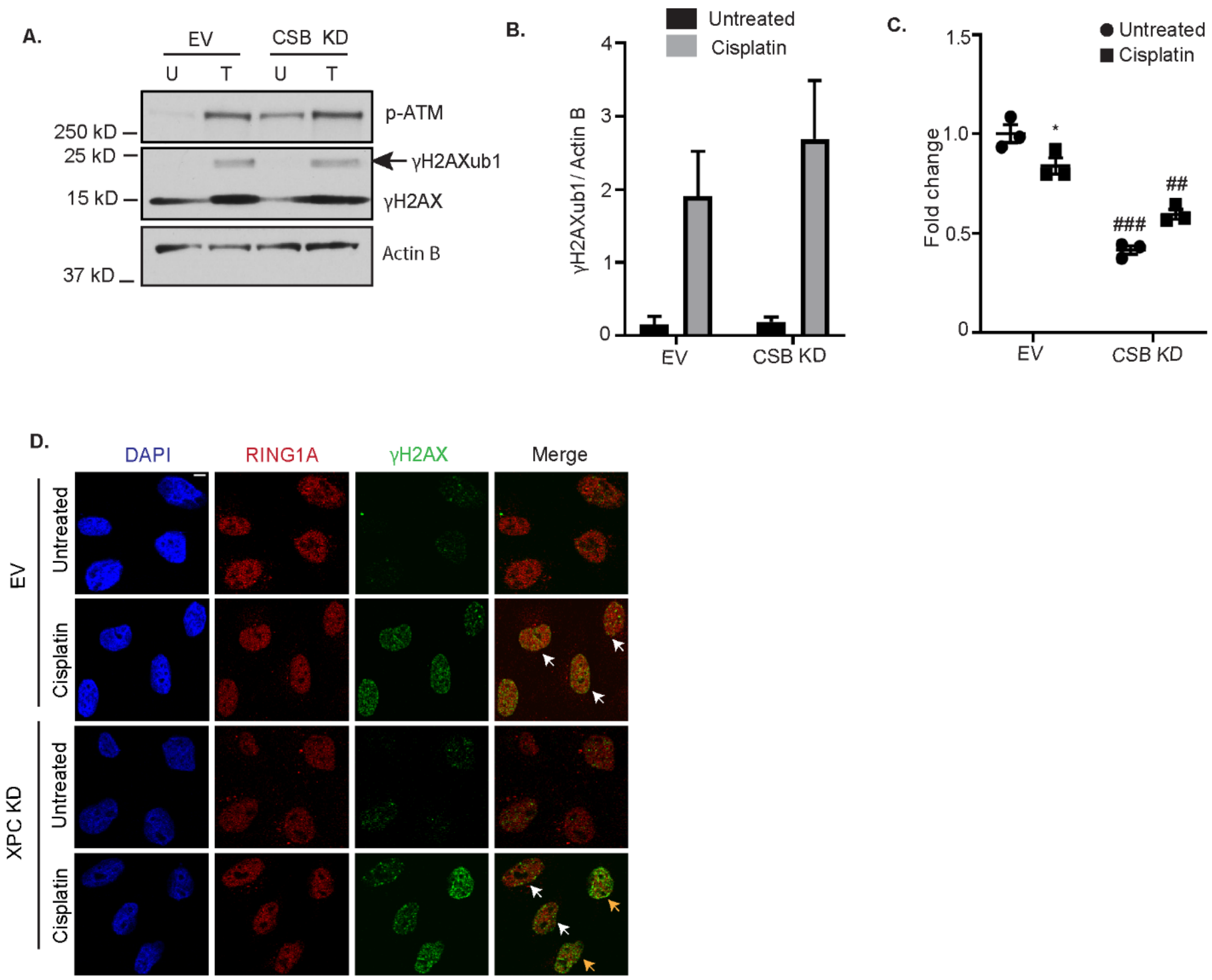


**Supplementary Figure S3: Effects of CSB or XPC knockdown on platinum-induced γH2AXub1**

**(A)** EV or CSB KD OVCAR5 cells were untreated (U) or treated with 12 μM cisplatin (T) for 8 hours. Lysates were analyzed by WB. **(B)** Graph shows mean densitometric analysis γH2AXub1 normalized to actin B ± SEM (N=3). **(C)** Graph shows mean CSB mRNA levels in untreated and cisplatin treated EV and CSB KD cells relative to EV untreated ± SEM (N=3). **(D)** Representative images of OVCAR5 EV and XPC KD cells showing RING1A colocalization with γH2AX. White arrows point to cells with colocalization and yellow arrows point to cells without colocalization (Note the decrease in yellow). Scale bar = 5 μM. Statistical significance was calculated using Student's t test. For untreated versus cisplatin, P-values * < 0.05, ** <0.005, *** < 0.0005. For EV versus CSB KD, P – values # < 0.05, ## < 0.005, ### < 0.005.

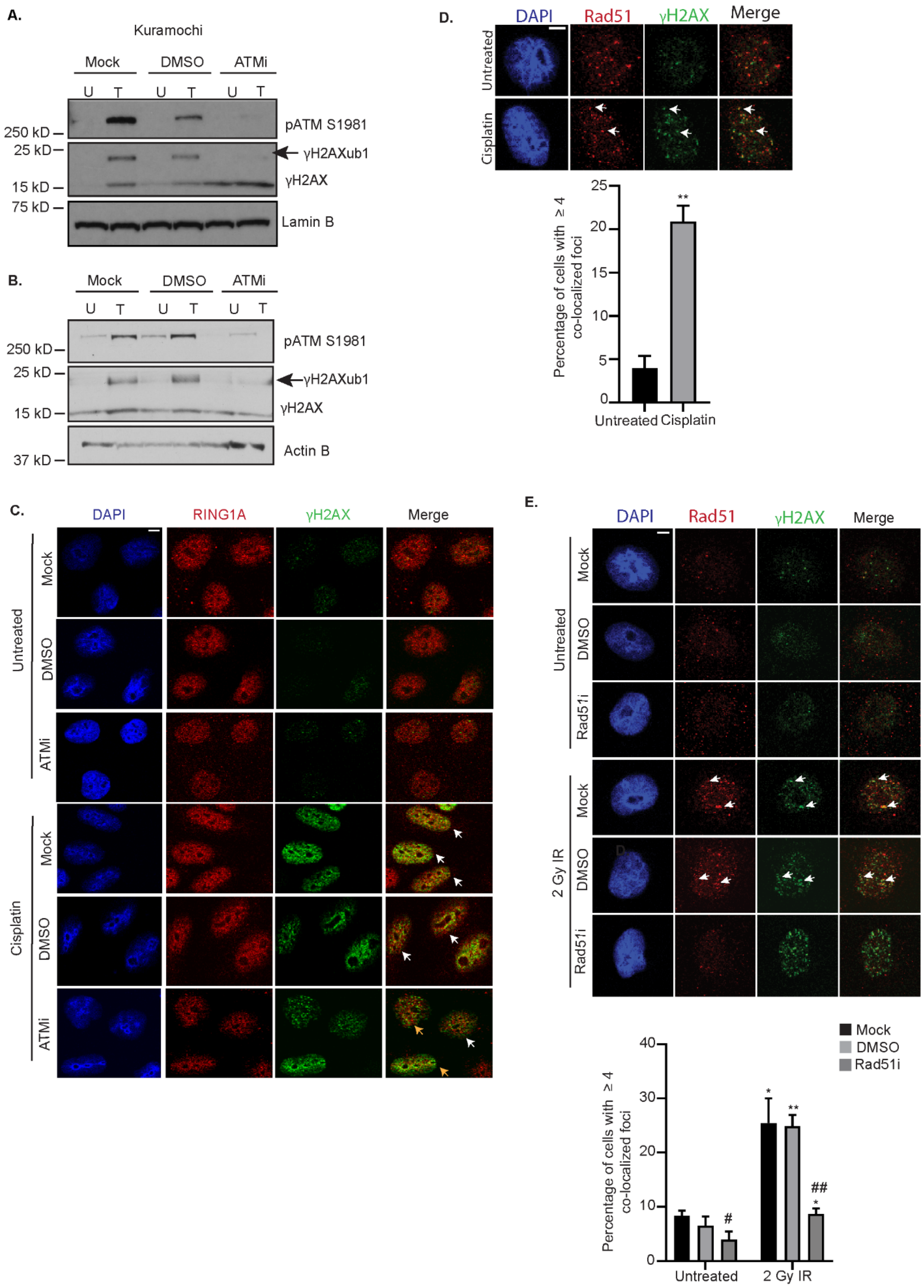
A.
Kuramochi
Mock
DMSO
ATMi
U T U T U T
250 kD
25 kD
15 kD
75 kD
pATM S1981
γH2AXub1
γH2AX
Lamin B
B.
Mock
DMSO
ATMi
U T U T U T
250 kD
25 kD
15 kD
37 kD
pATM S1981
γH2AXub1
γH2AX
Actin B
C.
DAPI
RING1A
γH2AX
Merge
Untreated
Cisplatin
Mock
DMSO
ATMi
D.
DAPI
Rad51
γH2AX
Merge
Untreated
Cisplatin
Percentage of cells with ≥ 4 co-localized foci
0
5
10
15
20
25
**
Untreated
Cisplatin
E.
DAPI
Rad51
γH2AX
Merge
Untreated
2 Gy IR
Mock
DMSO
Rad51i
Percentage of cells with ≥ 4 co-localized foci
0
10
20
30
40
#
*
**
##
Untreated
2 Gy IR

**Supplementary Figure S4: Rad51 colocalizes with γH2AX in response to cisplatin treatment**

**(A)** Kuramochi cells were not pretreated (mock) or pretreated with DMSO or 15 µM ATM inhibitor Ku-55933 for 1 hour and then untreated (U) or treatment with 15 µM cisplatin for 8 hours (T) (N=3). **(B)** OVCAR5 cells were pretreated with ATM inhibitor as in (A) and then untreated (U) or treated with 6.77 µM carboplatin for 24 hours. Lysates were analyzed by WB (N=2). **(C)** Representative images of OVCAR5 cells showing RING1A colocalization with γH2AX with and without ATMi followed by cisplatin treatment. White arrows indicate cells showing colocalization and yellow arrows indicate cells with less or no colocalization. **(D)** OVCAR5 cells were either untreated or treated with 12 µM cisplatin for 8 hours and then immunofluorescence was performed for γH2AX (green) and Rad51 (red). White arrows point to examples of γH2AX and Rad51 foci that co-localize. Graph shows mean percentage of cells with ≥ 4 Rad51 and γH2AX co-localized foci ± SEM. Scale bar = 5 µm. **(E)** OVCAR5 cells were not pretreated (mock) or pre-treated with DMSO or 50 µM Rad51 inhibitor B02 for 2 hours and then untreated (U) or treated with 2 Gy IR. After IR exposure, cells were allowed to recover for 15 minutes at 37$^{o}$C. Graph shows mean percentage of cells with ≥ 4 γH2AX and Rad51 co-localized foci ± SEM (N=3). White arrows show examples of γH2AX and Rad51 foci that co-localize. Scale bar = 5 µM. Statistical significance was calculated using Student's t test. For untreated versus cisplatin or 2 Gy IR treated, P-values * < 0.05, ** <0.005, *** < 0.0005. For Mock or DMSO versus Rad51i P – values # < 0.05, ## < 0.005, ### < 0.005.

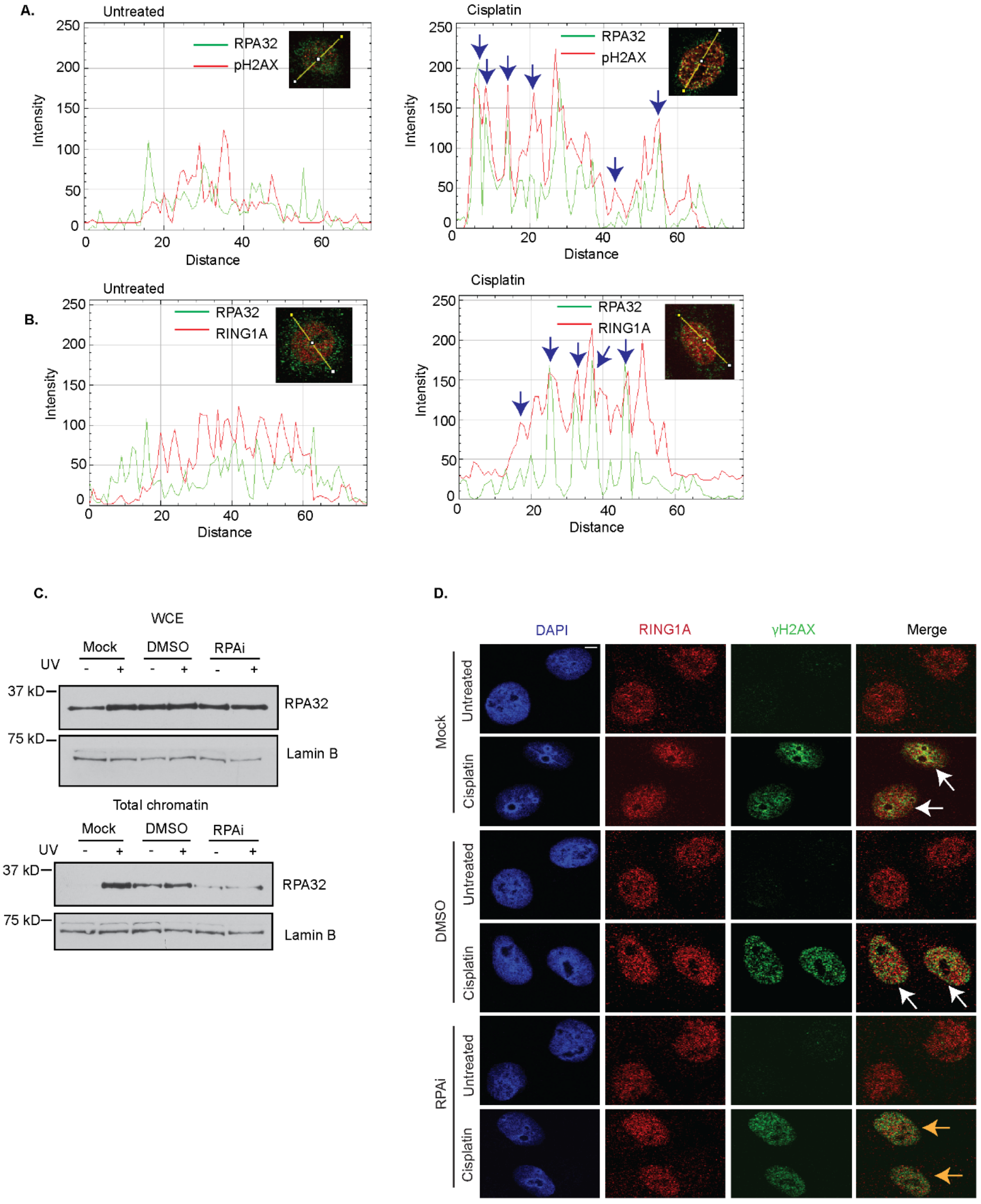
A.
Untreated
RPA32
pH2AX
Intensity
Distance
Cisplatin
B.
RING1A
C.
WCE
Mock
DMSO
RPAi
UV
37 kD
75 kD
Lamin B
Total chromatin
D.
DAPI
RING1A
γH2AX
Merge
Untreated
Cisplatin

**Supplementary Figure S5: RPA inhibition reduces RING1A localization to sites of cisplatin-induced DNA damage**

**(A)** Representative RGB profile of untreated and cisplatin treated cell from Figure 4B showing RPA32 colocalization with γH2AX. Blue arrows indicate foci which colocalize. **(B)** Representative RGB profile of untreated and cisplatin treated cell from Figure 4C showing RPA32 colocalization with RING1A. Blue arrows show colocalizing foci. **(C)** OVCAR5 cells were not pre-treated (Mock) or pre-treated with the vehicle (DMSO) or with 8 μM RPA inhibitor NERx329 (RPAi) and then untreated (-) or treated with 200 $mJ/cm^2$ UV (+) followed by 15 minutes recovery. Then total chromatin extraction was performed and lysates were analyzed by western blot. WCE – whole cell extract (N =3). **(D)** Representative images of RING1A and γH2AX colocalization in mock/DMSO/RPAi pretreated followed by untreated and cisplatin treated OVCAR5 cells. White arrows indicate cells with colocalization while yellow arrows indicate cells with less colocalization (Note the decrease in yellow color foci).

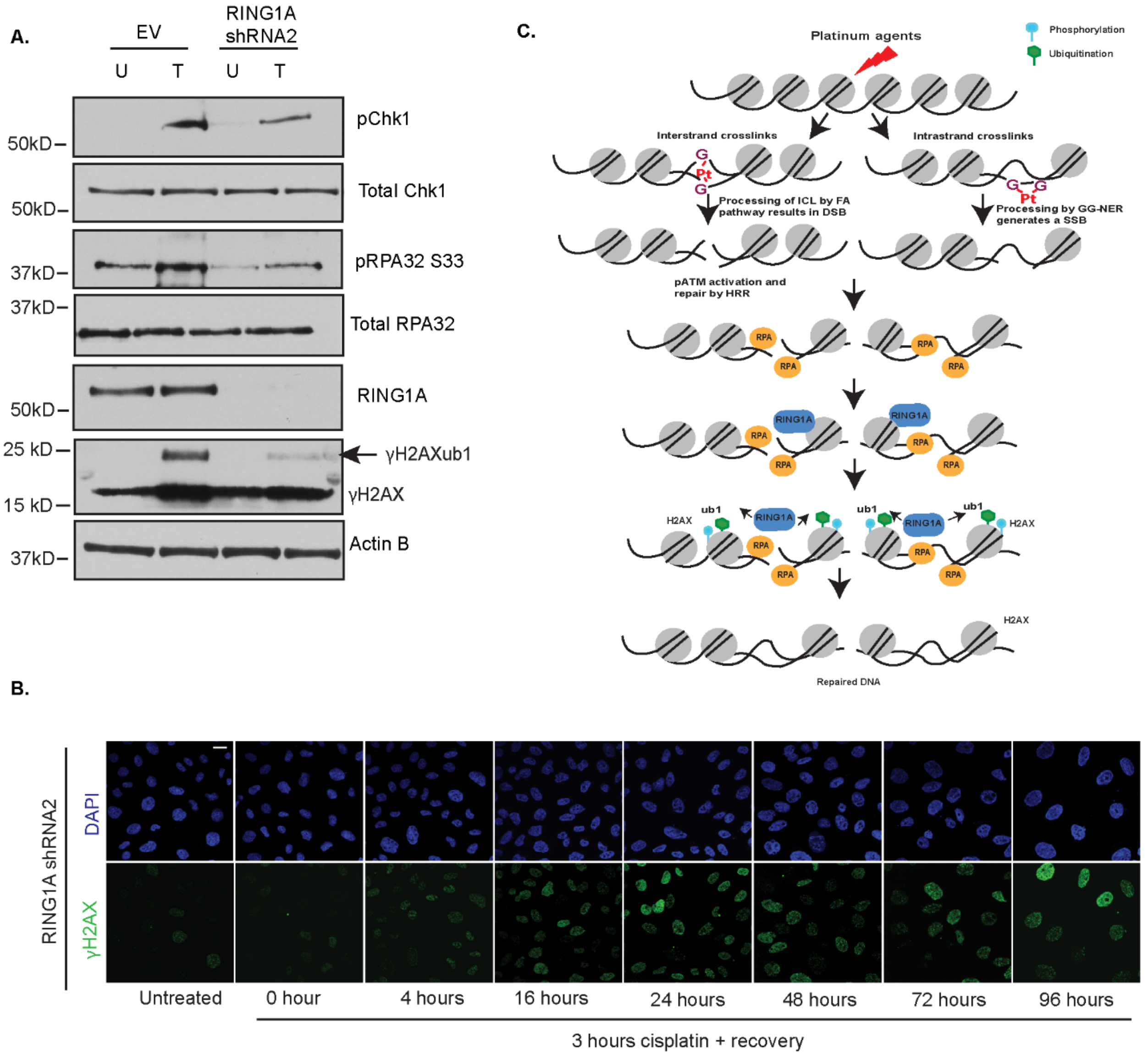


**Supplementary Figure S6: RING1A knockdown reduces phosphorylation of pS345 Chk1 and repair of cisplatin DNA damage**

**(A)** OVCAR5 cells infected with EV or RING1A viral shRNA2 were untreated and treated with cisplatin for 8 hours. Cell lysates were collected and analyzed by WB. **(B)** OVCAR5 cells infected RING1A viral shRNA2 were treated with 6μM cisplatin and allowed to recover for the indicated time points. Representative images for γH2AX at the indicated time points are shown. Scale bar = 20 μm. **(C)** Model for platinum-induced γH2AXub1 in OC cells. Treatment of OC cells with platinum agents results in formation of both ICLs and intrastrand crosslinks. ICLs are processed by the FA pathway resulting in a DSB while intrastrand crosslinks are processed by the GG-NER pathway generating a SSB. Resulting DSBs activate ATM and are repaired by HRR while SSBs are bound by RPA and repaired by GG-NER. RPA functioning in both HRR and GG-NER pathways binds to ssDNA and mediates RING1A localization to sites of platinum-induced DNA damage. RING1A contributes to the monoubiquitination of γH2AX facilitating the repair of damage.
